# Expanded genetic landscape of chronic obstructive pulmonary disease reveals heterogeneous cell type and phenotype associations

**DOI:** 10.1101/355644

**Authors:** Phuwanat Sakornsakolpat, Dmitry Prokopenko, Maxime Lamontagne, Nicola F. Reeve, Anna L. Guyatt, Victoria E. Jackson, Nick Shrine, Dandi Qiao, Traci M. Bartz, Deog Kyeom Kim, Mi Kyeong Lee, Jeanne C. Latourelle, Xingnan Li, Jarrett D. Morrow, Ma’en Obeidat, Annah B. Wyss, Xiaobo Zhou, Per Bakke, R Graham Barr, Terri H. Beaty, Steven A. Belinsky, Guy G. Brusselle, James D. Crapo, Kim de Jong, Dawn L. DeMeo, Tasha E. Fingerlin, Sina A. Gharib, Amund Gulsvik, Ian P. Hall, John E. Hokanson, Woo Jin Kim, David A. Lomas, Stephanie J. London, Deborah A. Meyers, George T. O’Connor, Stephen I. Rennard, David A. Schwartz, Pawel Sliwinski, David Sparrow, David P. Strachan, Ruth Tal-Singer, Yohannes Tesfaigzi, Jørgen Vestbo, Judith M. Vonk, Jae-Joon Yim, Yohan Bossé, Ani Manichaikul, Lies Lahousse, Edwin K. Silverman, H. Marike Boezen, Louise V. Wain, Martin D. Tobin, Brian D. Hobbs, Michael H. Cho, International COPD Genetics Consortium

## Abstract

Chronic obstructive pulmonary disease (COPD) is the leading cause of respiratory mortality worldwide. Genetic risk loci provide novel insights into disease pathogenesis. To broaden COPD genetic risk loci discovery and identify cell type and phenotype associations we performed a genome-wide association study in 35,735 cases and 222,076 controls from the UK Biobank and additional studies from the International COPD Genetics Consortium. We identified 82 loci with P value < 5×10^−8^; 47 were previously described in association with either COPD or population-based lung function. Of the remaining 35 novel loci, 13 were associated with lung function in 79,055 individuals from the SpiroMeta consortium. Using gene expression and regulation data, we identified enrichment for loci in lung tissue, smooth muscle and alveolar type II cells. We found 9 shared genomic regions between COPD and asthma and 5 between COPD and pulmonary fibrosis. COPD genetic risk loci clustered into groups of quantitative imaging features and comorbidity associations. Our analyses provide further support to the genetic susceptibility and heterogeneity of COPD.

## Background

Chronic obstructive pulmonary disease (COPD) is a disease of enormous and growing global burden^1^, ranked third as a global cause of death by the World Health Organization in 2016^2^. Environmental risk factors, predominately cigarette smoking, account for a large fraction of disease risk, but there is considerable variability in COPD susceptibility among individuals with similar smoking exposure. Studies in families and in populations demonstrate that genetic factors account for a substantial fraction of disease susceptibility. Similar to other adult-onset complex diseases, common variants likely account for the majority of population risk^3,4^. Our previous efforts identified 22 genome-wide significant loci^5^. Expanding the number of risk loci can lead to novel disease pathogenesis insights not only through discovery of novel biology^6,7^ but also through informing more global insights such as functional links between loci and cell-type and phenotype identification driving COPD genetic risk^5^.

We performed a genome-wide association study including previously described studies from the International COPD Genetics Consortium (ICGC) with additional subjects from UK Biobank^8^, a population-based study of several hundred thousand subjects with lung function and cigarette smoking assessment. We determined, through bioinformatic and computational analysis, the likely set of variants, genes, cell types, and biologic pathways implicated by these associations. Finally, we assessed our genetic findings for relevance to COPD-specific, respiratory, and other phenotypes.

## Results

### Genome-wide association study of COPD

We included a total of 257,811 individuals from 25 studies in the analysis, including studies from International COPD Genetics Consortium and UK Biobank. We defined COPD based on pre-bronchodilator spirometry using pre-bronchodilator spirometry according to modified Global Initiative for Chronic Obstructive Lung Disease (GOLD) criteria for moderate to very severe airflow limitation^9^, resulting in 35,735 cases and 222,076 controls (**Supplementary Tables See** the Excel file.

Supplementary Table 1). We tested association of COPD and 6,224,355 variants in a meta-analysis of 25 studies using a fixed-effects model. We found no evidence of confounding by population substructure using linkage disequilibrium score regression (LDSC) intercept (1·0377, s.e. 0.0094).

We identified 82 loci (defined using 2-Mb windows) at genome-wide significance (P < 5 × 10^−8^) (**Figure 2**).Forty-seven of 82 loci were previously described as genome-wide significant in COPD^5,10–12^ or lung function^13,14,23,15–22^ (**Supplementary Table 2**), leaving 35 novel loci (**Table 1**). We then sought to replicate these novel loci. Given the strong genetic correlation between population-based lung function and COPD, we tested the lead variant at each for association with FEV_1_ or FEV_1_/FVC in 79,055 individuals from SpiroMeta. We identified 13 loci - *C1orf87*, *DENND2D*, *DDX1*, *SLMAP*, *BTC*, *FGF18*, *CITED2*, *ITGB8*, *STN1*, *ARNTL*, *SERP2*, *DTWD1*, and *ADAMTSL3* that replicated using a Bonferroni correction for a one-sided P < 0·05/35; **Table 1**). Although not meeting the strict Bonferroni threshold, additional 14 novel loci were nominally significant in SpiroMeta (consistent direction of effect and one-sided P < 0·05): *ASAP2*, *EML4*, *VGLL4*, *ADCY5*, *HSPA4*, *CCDC69*, *RREB1*, *ID4*, *IER3*, *RFX6*, *MFHAS1*, *COL15A1*, *TEPP*, and *THRA* (**Table 1**).In our overall meta-analysis, all 82 of the genome-wide significant loci showed consistent direction of effect with either FEV_1_ or FEV_1_/FVC ratio in SpiroMeta (**Table 1** and **Supplementary Table 2**). We note that 9 of our 35 novel loci were recently described in a contemporaneous analysis of lung function in UK Biobank^24^. None of the novel loci appeared to be due to cigarette smoking (**Supplementary Results**). Including all 82 genome-wide significant variants, we explain up to 7·0 % of the phenotypic variance in liability scale, using a 10% prevalence of COPD, and acknowledging that these effects are likely overestimated in the discovery sample. This represents a 48% increase in COPD phenotypic variance explained by genetic loci compared to the 4·7% explained by 22 loci reported in a recent GWAS of COPD^5^.

**Figure 1.**
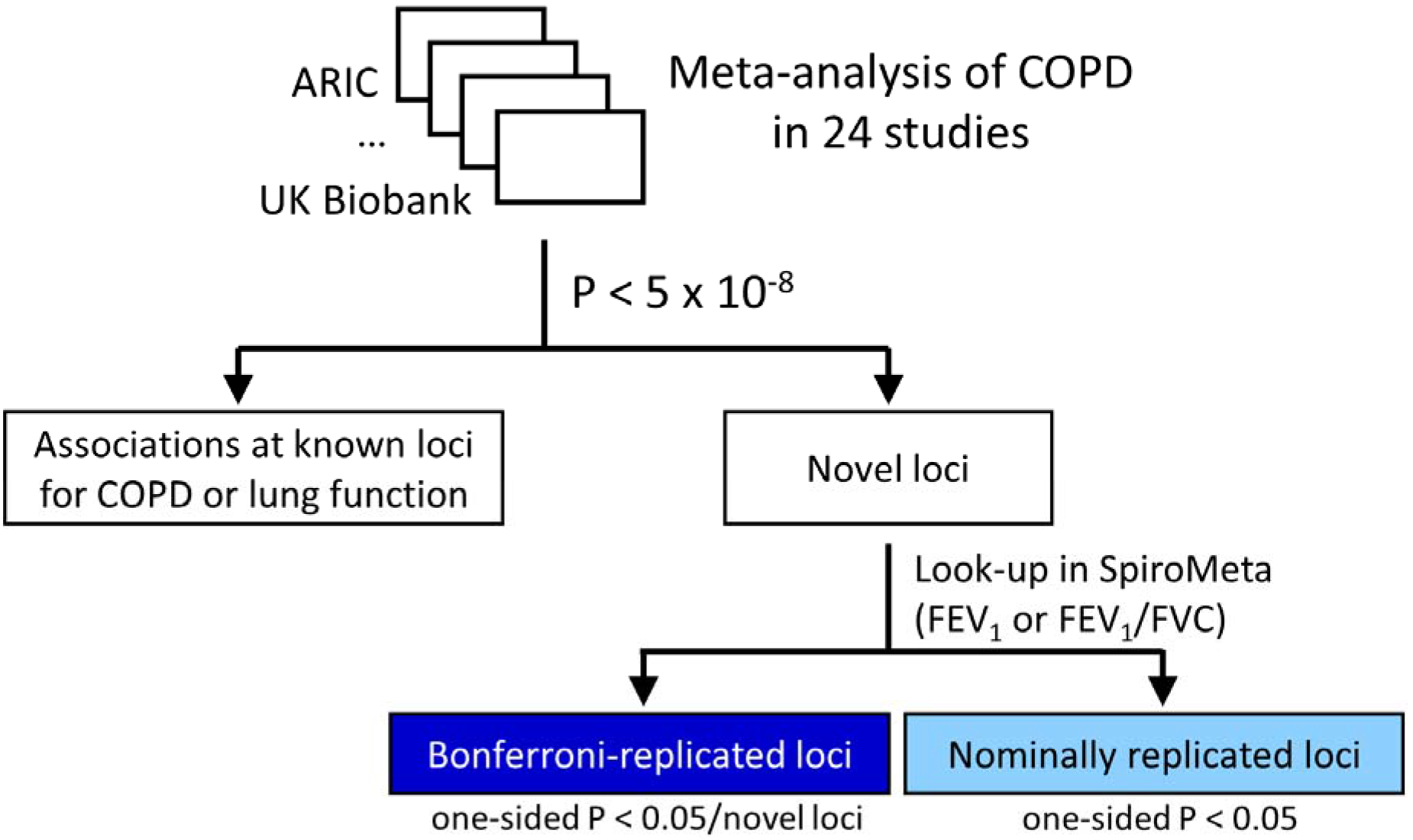
Study design. COPD, chronic obstructive pulmonary disease; FEV_1_, force expiratory volume in one second; FVC, forced vital capacity.

**Figure 2.**
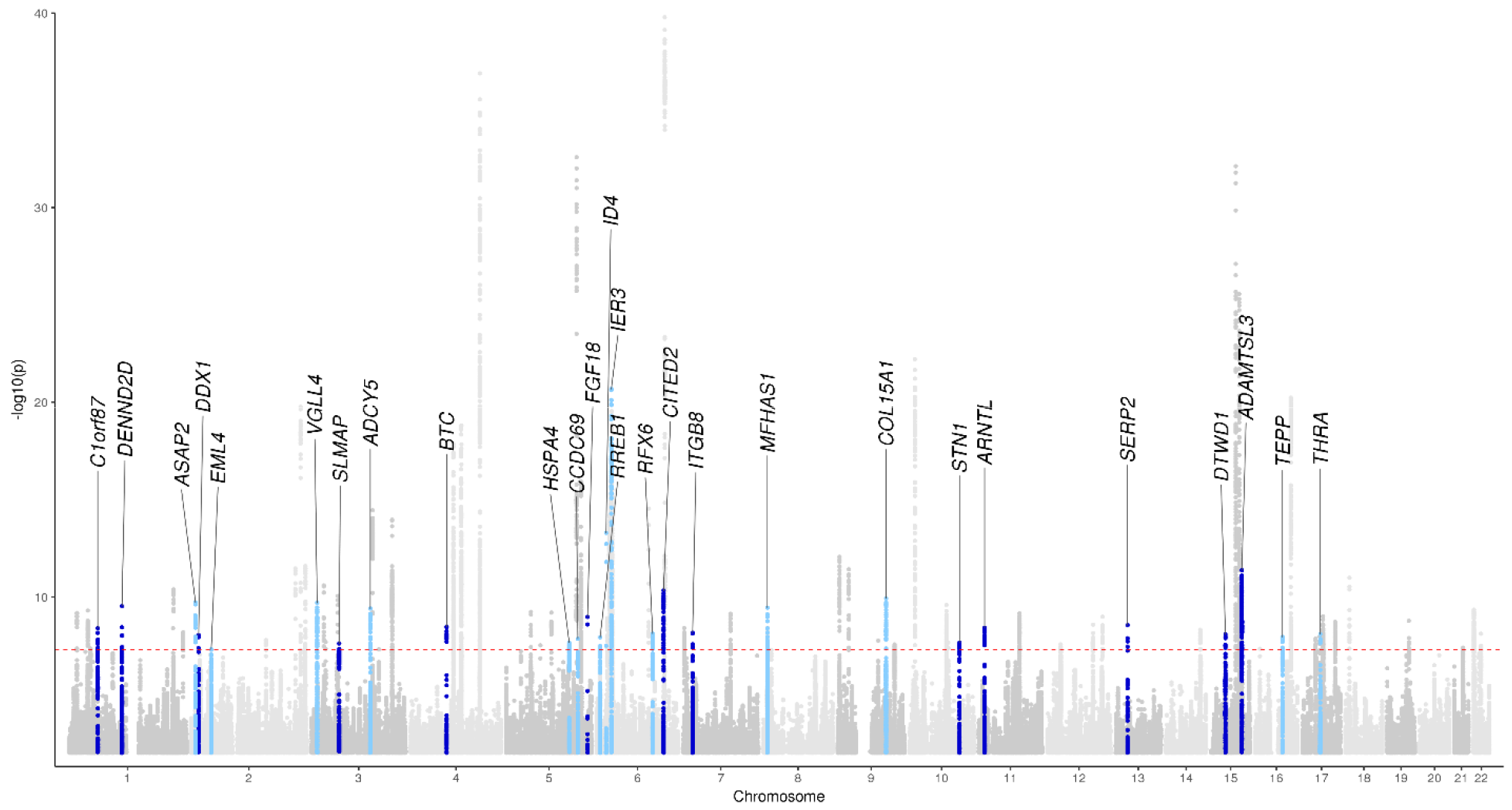
Manhattan plot. Loci are labeled with the closest gene to the lead variant. Colors indicates variants at novel loci which replicated using Bonferroni-corrected threshold in SpiroMeta (dark blue, one sided P < 0.05/35) and nominally significant threshold (light blue, one-sided P < 0.05).

### Identification of secondary association signals

We then used approximate conditional and joint analysis to find secondary signals at each of the 82 genome-wide significant loci. We found 82 secondary signals at 50 loci, resulting in a total of 164 independent associations in 82 loci (**Supplementary Table 3**). Of 50 loci containing secondary associations, 33 were at loci previously described for COPD or lung function, and 6 at Bonferroni-replicated novel loci. Of 82 secondary associations, 20 reached genome-wide significance (P < 5 × 10^−8^) (**Supplementary Table 3**). Of 61 novel (not previously described in COPD or lung function) independent associations, 21 reached a region-wise Bonferroni-corrected threshold (one-sided P < 0.05/novel independent association(s) in each locus) in unconditioned associations from SpiroMeta (**Methods**and **Supplementary Table 3**).

### Tissue and specific cell types

In determining the tissue in which COPD genetic variants function to increase COPD risk, lung is the obvious tissue to consider; however, COPD is a systemic disease^25,26^ and within the lung, the specific cell-types underlying disease pathogenesis are largely unknown. Furthermore, available databases often include cell types (e.g. smooth muscle) from non-lung organs (e.g. the gastrointestinal tract). To identify putative causal tissues and cell types, we assessed the enrichment of our genome-wide significant COPD loci in integrated genome annotations at the single tissue level^27^, tissue-specific epigenomic marks^28^, and genome-wide gene expression patterns^29^. Lung tissue showed the most significant enrichment (OR 9·25, P=1·36 × 10^−9^), as previously described, though significant enrichment was also seen in heart (OR 6·85, P=3·83 × 10^−8^) and the gastrointestinal (GI) tract (OR 5·53, P=6·45 × 10^−11^). In an analysis of enriched epigenomic marks, the most significant enrichment was in fetal lung and GI smooth muscle DNase hypersensitivity sites (DHS) (P= 6·75 × 10^−8^) and H3K4me1 (P= 7·31 × 10^7^) (**Supplementary Table 4**). To further identify lung-specific cell types, we tested whether 47 known and 13 novel COPD-associated loci contain genes specifically expressed in data sets from single-cell RNA-seq. We found enrichment of alveolar type II and basal cells (P=0·016 and P=0·042, respectively) using single-cell RNA-seq gene expression data from respiratory cell types^30^ (**Supplementary Figure 3** and **Supplementary Table 5** for individual locus results).

### Fine-mapping of associated loci

To identify the most likely causal variants, we performed fine mapping using Bayesian credible sets^31^. Including 160 potential primary and secondary association signals (excluding four variants in the major histocompatibility complex (MHC) region), 61 independent signals had a 99% credible set with fewer than 50 variants; 34 signals had credible sets with fewer than 20 variants (**Supplementary Figure 4**). Only the association signal at *NPNT* (4q24) could be fine-mapped to single variant; however, in 17 other loci, a single variant had posterior probability of driving association (PPA) greater than 60% (**Supplementary Table 6**). Most sets included variants that overlapped genic enhancers of llung-related cell types (e.g., fetal lung fibroblasts, fetal lung, and adult lung fibroblasts) and were predicted to alter transcription binding motifs (**Supplementary Table 6**). Of 61 credible sets with fewer than 50 variants, eight sets contained at least one deleterious variant. These deleterious variants included 1) missense variants affecting *TNS1*, *RIN3*, *GPR126*, *ADAM19*, *ATP13A2*, *BTC*, and *CRLF3*; and 2) a splice donor variant affecting a lincRNA - AP003059.2.

### Candidate target genes

In most cases, the closest gene to a lead SNP will not be the gene most likely to be the causal or effector gene of disease-associated variants^32–34^. Thus, to identify the potential effector (‘target’) genes underlying these genetic associations, we integrated additional molecular information including gene expression; gene regulation (open chromatin and methylation data), chromatin interaction, co-regulation of gene expression with gene sets and coding variant data (**Methods** and **Figure 3a**).

**Figure 3.**
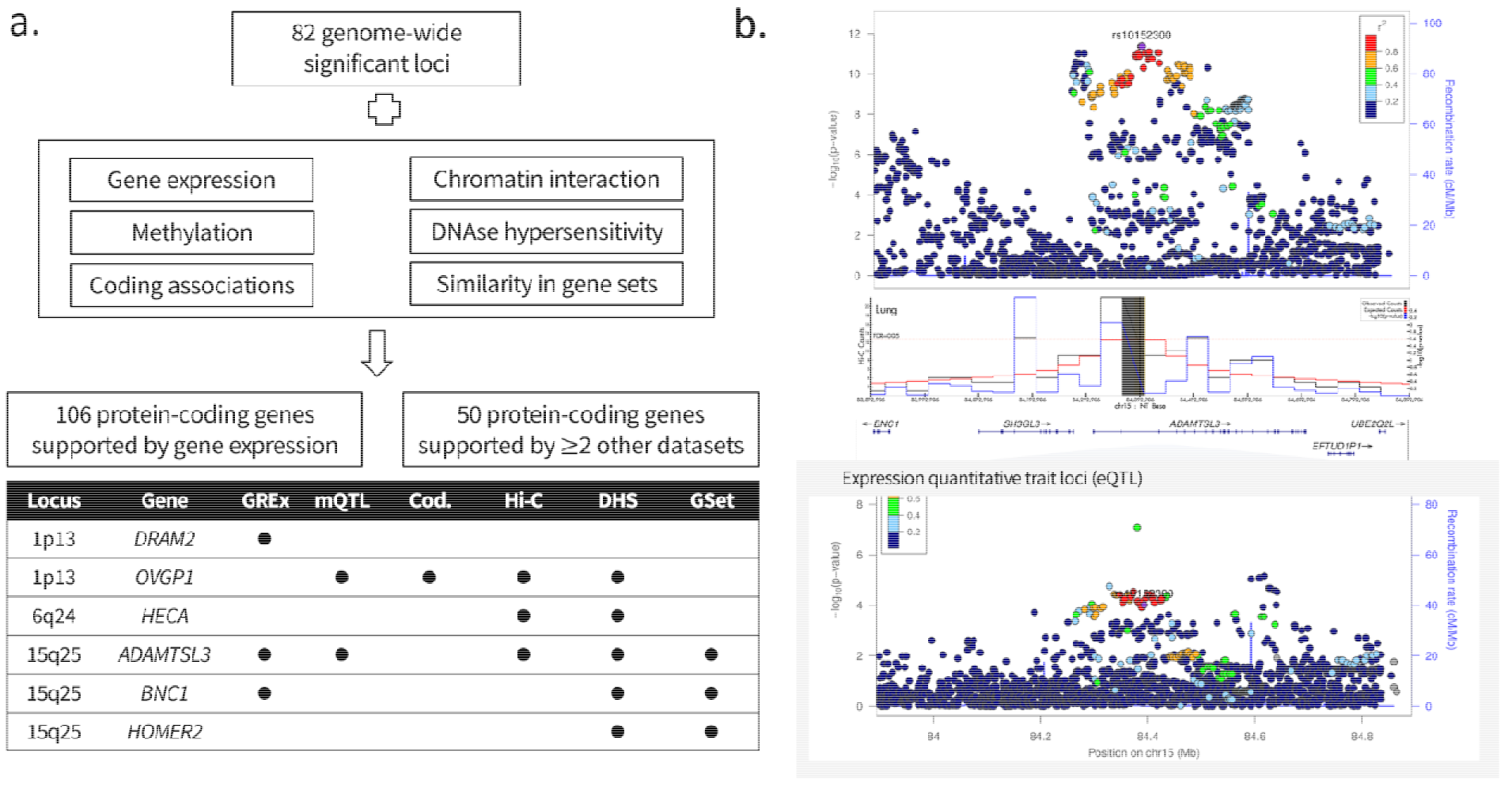
Identification of target genes. (a) Overview of datasets used to identify target genes at genome-wide significant loci (b) Regional association plots at *ADAMT5L3* locus showing GWAS (top), chromatin interaction in lung tissue (middle) and expression quantitative trait loci (bottom).e.

At 82 loci, 472 genes were implicated by analysis of least one dataset; 106 genes were implicated by gene expression (Bonferroni corrected at locus level), and an additional 50 genes by >= 2 other datasets (methylation, chromatin interaction, open chromatin regions, similarity in gene sets or deleterious coding variants (**Figure 3a**)),for a total of 156 genes meeting more strigent criteria. Excluding loci in the extended MHC region, the median numbers of potentially implicated genes per locus was four with a maximum of 17 genes (7q22.1 and 17q21.1). The median distance of implicated genes to top associated variants was 346 Kb, restricting to genes +/−1 Mb of top associated variants. Among 82 loci, 60 loci (73%) included the nearest gene. We identified 20 genes which were region-wise Bonferroni significant in exome sequencing data. Two genes (*ADAM19* and *ADAMTSL3*) were implicated by five datasets (**Figure 3b**)and another two (*EML4* and *RIN3*) were implicated by four datasets. A summary of all genes implicated using these approaches in **SupplementaryTable 7**.

### Associated pathways

To gain further functional insight of associated genetic loci, we performed gene-set enrichment analysis using DEPICT. Among 165 enriched gene sets at FDR < 5%, 44% of them were related to the developmental process term, with lung development P = 1.02 × 10^−6^; significant sub-terms included lung alveolus development (nominal P= 0·0003) and lung morphogenesis (nominal P= 0·0005). We also found enrichment of extracellular matrix-related pathways including laminin binding, integrin binding, mesenchyme development, cell-matrix adhesion, and actin filament bundles. Additional pathways of note included histone deacetylase binding, Wnt receptor signaling pathway, SMAD binding, the MAPK cascade, and transmembrane receptor protein serine/threonine kinase signaling pathway. Full enrichment analysis results including the top genes for each DEPICT gene set are shown in **Supplementary Table 8**.

### Phenotypic effects of known and novel associations for COPD

To characterize the phenotypic effects of 82 genome-wide significant loci, we performed a phenome-wide association analysis within the deeply phenotyped COPDGene study^35^. We tested the overall structure of associated phenotypes by using hierarchical clustering across scaled Z scores of associations. We identified two clusters of associated variants (**Supplementary Figure 5**). As these two clusters appeared to predominantly differentiate among imaging features, we repeated variant clustering limited to quantitative computed tomography (CT) imaging features. We found two clusters of variants, differentiated by association with quantitative emphysema, emphysema distribution, gas trapping, and airway phenotypes (**Figure 4a**). We then evaluated the association of the 82 genome-wide significant variants in a prior GWAS of emphysema and airway quantitative CT features^36^ (**Supplementary Table 10**).

**Figure 4.**
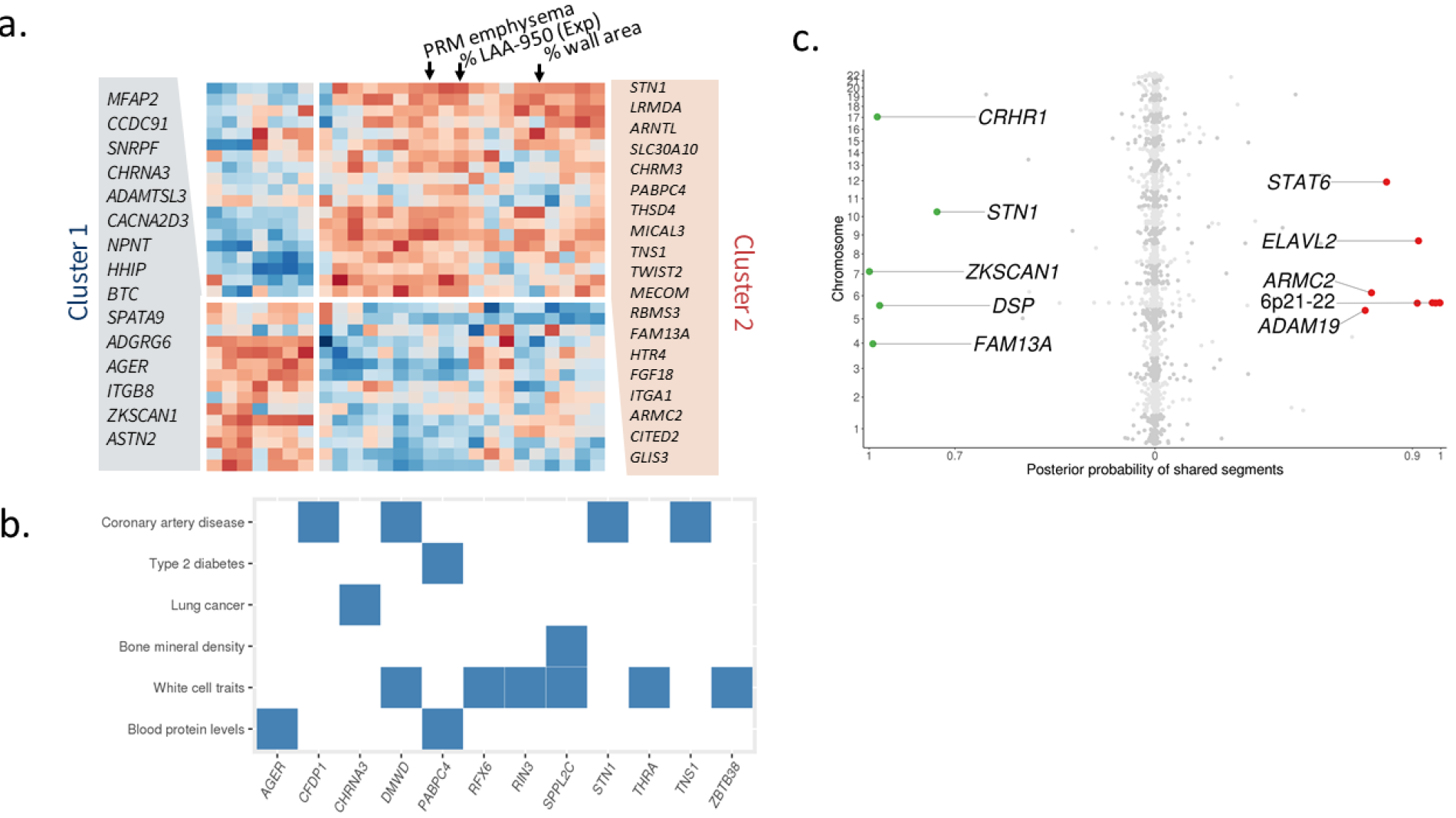
Effects on COPD-related and other phenotypes. (a) Heatmap of associations of 60 genome-wide significant variants (known and replicated novel associations) and imaging phenotypes in COPDGene (b) Overlapping of genome-wide significant loci of COPD and select traits from GWAS Catalog (c) Genome-wide overlapping results between COPD with pulmonary fibrosis (left) and asthma (right). PRM emphysema, emphysema quantified by parametric response mapping; %LAA-950 (Exp), percentage of low attenuation area at-950 Hounsfield Units at expiration (related to gas trapping). GREx, Genetically regulated gene expresssion (significant associations identified by S-PrediXcan); mQTL, methylation quantitative trait loci (colocalized signals between GWAS and mQTL at posterior probability > 04); Cod., Coding associations (significant single variant or gene-based association tests for deleterious coding variants); Hi-C, significant chromatin interaction identified in human lung or IMR90 cell line; DHS, DNase hypersensititvity sites (using regulatory fine-mapping or regfm software); GSet, target genes identified by DEPICT using reconstituted gene sets.

We also examined all genome-wide significant loci in the NHGRI-EBI GWAS Catalog^37^ (**Supplementary Figure 6 and Supplementary Table 11**) and looked for variants in linkage disequilibrium (r^2^ > 0·2) with the lead GWAS variant. Many variants associated with anthropometric measures including height and body mass index (BMI), measurements on blood cells (red and white cells), and cancers. COPD is well known for having common comorbidities, such as coronary artery disease (CAD), type 2 diabetes mellitus (T2D), bone density, and lung cancer. Of these diseases, we only found evidence of modest overall genetic correlation (using linkage disequilibrium score regression) between COPD and lung cancer (**Supplementary Results**). However, at individual loci, and using more stringent linkage disequilibrium (r^2^ > 0·6), we found evidence of shared risk factors for COPD, including a genome-wide significant variant near *PABPC4* associated with T2D, four variants with CAD (near *CFDP1*, *DMWD*, *STN1*, and *TNS1*), and a variant near *SPPL2C* with bone density (**Figure 4b**).

### Identification of loci overlapping with asthma and pulmonary fibrosis

Based on our previous identification of genetic overlap of COPD with asthma, and COPD with pulmonary fibrosis, we also examined loci for specific overlap with these two diseases. In asthma, we noted an r^2^ > 0.2 with one of our variants, and previously reported variants and *ID2*, *ZBTB38*, *C5orf56*, *MICA*, *AGER*, *HLA-DQB1*, *ITGB8*, *CLEC16A*, and *THRA*. In pulmonary fibrosis, in addition to our previously described overlap between *FAM13A*, *DSP*, and *17q21*, we noted *ZKSCAN1* and *STN1* (**Supplementary Table 11**).To more closely examine overlap, applied a Bayesian method (gwas-pw) of COPD associations from our current GWAS with previous GWASs of asthma (limited to those of European ancestry) and pulmonary fibrosis^38,39^. To mitigate the effect of including asthma cases in the GWAS of COPD, we excluded individuals with self-reported asthma from the UK Biobank (**Methods** and **Supplementary Results**).We identified 14 shared genome segments (posterior probability > 70%), nine with asthma and five with pulmonary fibrosis (**Figure 4c** and **Supplementary Table 9**).Of nine segments shared with asthma, five segments reside within the MHC region (6p21-22). Non-MHC segments included loci near *ADAM19*, *ARMC2*, *ELAVL2*, and *STAT6*. Of five segments shared with pulmonary fibrosis, two segments were identified including loci near *ZKSCAN1*, *STN1* (formerly known as *OBFC1*), in addition to the three segments identified in the previous study^5^ (*FAM13A*, *DSP*, and the 17q21 locus, here *CRHR1*). For all overlapping loci between COPD and asthma, overlapping variants had the same direction of effect (i.e., increasing risk for both COPD and asthma). Conversely, shared variants between COPD and pulmonary fibrosis all had an opposite effect (i.e., increasing risk for COPD but protective for pulmonary fibrosis).

## Discussion

Genetic factors play an important role in COPD susceptibility. We examined genetic risk of COPD in a genome-wide association study including a total of 35,735 cases and 222,076 controls. We identified 82 genome-wide significant loci for COPD, of which 47 were previously identified in genome-wide association studies of COPD, or population-based lung function. Of 35 loci not previously described, 13 replicated in an independent study of population-based lung function. Our results identify important disease pathways and may further explain the clinical heterogeneity seen in COPD.

Our study supports the role of early life events in the risk of COPD. Gene set enrichment analysis on our putative causal genes identified lung morphogenesis and lung alveolar development, the canonical Wnt receptor signaling pathway^40^^41^, the MAPK cascade, Ras protein signal transduction, and the nerve growth factor receptor signaling pathway. Further, the importance of gene regulation at fetal stages was highlighted through enrichment of heritability in epigenomic marks of various fetal tissues, with fetal lung showed the strongest signals. Our findings are consistent with recent epidemiologic studies demonstrating that a substantial portion of the risk of COPD may be develop in early life: genetic variants may set initial lung function^42^ and patterns of growth^42–44^.

We also identified several genes and pathways of interest not primarily related to lung development, some of which have been previously identified in studies of lung function^13^, including mesenchyme development and extracellular matrix, cilia structure, elastin-associated microfibrils, and retinoic acid receptor beta^45–47^. We used several data sources to attempt to assign causal gene at each locus, identifying 156 genes at 82 loci that were supported by either gene expression or a combination of at least 2 other data sources. One of our genes with the most support was *ADAMTSL3*. In addition to a role in development, this gene plays a role in cell-matrix interactions or in assembly of specific extracellular matrices^48^. Another novel finding was an association with the chitinase acidic (*CHIA*) gene at 1p13.3, which encodes a protein that degrades chitin^49^ and exhibits lung-specific expression^50,51^. *CHIA* variants have been associated with FEV_1_^52^, asthma^53–56^, and acid mammalian chitinase activity^55,57^. Its role in airway inflammation was demonstrated in an animal model of asthma^58^. Interleukin 17 receptor D (*IL17RD*) at 3p14.3 encodes a membrane protein belonging to the interleukin-17 receptor (IL-17R) protein family^49^. The gene product affects fibroblast growth factor signaling, inhibiting or stimulating growth through MAPK/ERK signaling^49^. It also interacts with TNF receptor 2 (TNFR2) to activate NF-κB^59^. Integrin subunit beta 8 (*ITGB8*) at 7p21.1 is a member of the integrin beta chain family and ITGB8 protein expression protein is increased in COPD^60–62^. This locus was also recently described in a separate study of allergic disease and asthma^63^. The *ITGB8* gene and encodes a single-pass type I membrane protein that binds to an alpha subunit to form an integrin complex^49^. The complex mediates cell-cell and cell-extracellular matrix interactions and plays a role in human airway epithelial proliferation^49^ and repair^64^.

In addition to identifying the effector gene, the effector cell type is of critical important for functional studies. We identified an overall enrichment of epigenomic marks in lung tissue and smooth muscle (also identified in studies of lung function^21^); this latter association is driven by non-respiratory cell types. Within a set of four lung cell types identified by single cell RNA-Seq, we identified enrichment for alveolar type 2 (AT2) cells, which recently have been shown to have regenerative properties^65^. The lung is comprised at least 40 different resident cell types^66^; thus, while our findings suggest cell types for further functional studies, they also highlight the need for profiling of additional lung cell types.

Characterization of genome-wide significant associations revealed heterogenous effect to COPD-related phenotypes and other biological processes. Within the well-phenotyped COPDGene cohort, we identified variable effects of these variants on computed tomography (CT) features, smoking status and intensity, diffusing capacity of carbon monoxide, asthma, and inflammatory biomarkers. Clustering these variants found differential effects on emphysema and airway phenotypes, a well-described source of heterogeneity in COPD^67–69^. Analyzing over hundreds of diseases/traits in GWAS Catalog, we identified overlapping associations with various diseases/traits in multiple organ systems, comorbidities such as coronary artery disease, bone mineral density, and type 2 diabetes mellitus. Together, the identification of variable COPD risk loci associations with sub-phenotypes and other diseases^70,71^ may have potential for more nuanced approaches to therapy for COPD.

We performed additional specific analysis in two diseases that overlap with COPD, asthma and pulmonary fibrosis. Extending from previously reported genetic correlations^5^, we identified 14 genetic loci shared between these pairs of diseases. Our analysis is the first to identify evidence for shared genetic segments between asthma and COPD. We identified multiple overlapping loci with asthma at the MHC region (6p21-22) and four other loci. The locus near *DDX1* was previously suggested to be associated with both COPD and asthma in a combined meta-analysis^72^, and is involved in NFkB pathway^73^, leading to a possible shared inflammatory mechanism between these diseases. In addition to three previously reported overlapping loci for COPD and IPF (*FAM13A* and *DSP*, both genome-wide significant, and 17q21, identified through Bayesian overlap analysis), we identified two loci near *ZKSCAN1* (7q22.1) and *STN1* (previously known as *OBFC1*, 10q24). The top associated variant near *STN1* is in linkage disequilibrium with a CAD-associated variant (rs12765878, r^2^=0·61). This locus has also been associated with leukocyte telomere length^74^, and lends additional support to a role of telomere maintenance as a common risk factor for these three diseases. Overall, our phenotype, gene-, and pathway- analyses illustrate the utility of both searching for enrichment of genetic signal overall and performing a more detailed identification of the effects of individual variants or groups of variants.

While our study is the largest genome-wide study of COPD, individuals meeting criteria for COPD in the UK Biobank may be different from other smoker-enriched studies, especially for smoking history. In addition, our use of population-based lung function for replication, along with pre-bronchodilator spirometry, could bias our findings against variants that are only associated with more severe forms of COPD. However, we observed concordant effect size estimates when including or excluding asthmatics. Thus, together with prior analyses^5^, our findings suggest that bias due to COPD case misclassification is likely small. However, we cannot rule out a role for studying more severe disease. We note that the alpha-1 locus (*SERPINA1*) was identified as genome-wide significant in smaller studies of emphysema and in severe COPD in smokers. In the current study, the association of the PIZ allele had P = 2·2 × 10^−5^ using moderate-to-severe cases, and a lower P-value (1·4 × 10^−6^) in severe (FEV_1_ < 50% predicted) cases despite a smaller sample size, an effect we have previously described^75^. Thus, despite the strong overlap of COPD with quantitative spirometry, new loci may be identified through studies of sufficiently large subsets of COPD patients with severe COPD or more specific and homogenous COPD phenotypes. Given suggestive evidence for replication using a related (but not identical) phenotype for additional novel loci beyond the 13 meeting Bonferroni, we chose to include all loci significant in discovery in subsequent analyses, recognizing that we likely included some false positive associations. Our study focused on relatively common variants, predominantly in individuals of European ancestry; more detailed studies of rare variants, HLA regions, and other ethnicities are warranted, but broader multi-ethnic analyses are limited by the number of cases in currently available cohorts.

The global burden of COPD is increasing. Our work finds a substantial number of new loci for COPD, and uses multiple lines of supportive evidence to identify potential genes and pathways for both existing and novel loci. Further investigation of genetic overlap and phenotypic effects finds new shared loci for asthma and idiopathic pulmonary fibrosis, and suggests heterogeneity across COPD-associated loci. Together, these insights provide multiple new avenues for investigation for this deadly disease.

## Methods

### Study populations

The UK Biobank is a population-based cohort consisting of 502,682 individuals^8^. To determine lung function, we used measures of forced expiratory volume in 1 second (FEV_1_) and forced vital capacity (FVC) derived from the spirometry blow volume-time series data, subjected to additional quality control based on ATS/ERS criteria^76^ (**Supplementary Methods**). We defined COPD in European-ancestry subjects using two pre-bronchodilator measurements of lung function according to modified Global Initiative for Chronic Obstructive Lung Disease (GOLD) criteria for moderate to very severe airflow limitation^9^: FEV_1_ less than 80% of predicted value (using reference equations from Hankinson et al.^77^), and the ratio between FEV_1_ and FVC less than 0.7. Genotyping was performed using Axiom UK BiLEVE array and Axiom Biobank array (Affymetrix, Santa Clara, California, USA) and imputed to the Haplotype Reference Consortium (HRC) version 1.1 panel^78^.

We invited participants in the prior International COPD Genetics Consortium (ICGC) COPD genome-wide association study to provide case-control association results (with the exception of the 1958 British Birth Cohort, to avoid overlapping samples with the replication sample). ICGC cohorts performed case-control association analysis based on pre-bronchodilator measurements of FEV_1_ and FEV_1_/FVC, and cases were identified using modified GOLD criteria, as above. Studies were imputed to 1000 Genomes reference panels. Detailed cohort descriptions and cohort-specific methods have been previously published^5^ (**Supplementary Methods**).

Based on the strong genetic overlap of lung function and COPD^5^, we performed lookups of select significant variants for FEV_1_ and FEV_1_/FVC in the SpiroMeta consortium meta-analysis^24^. Briefly, SpiroMeta comprised of a total of 79,055 individuals from 22 studies imputed to either the 1000 Genomes Project Phase 1 reference panel (13 studies) or the HRC (9 studies). Each study performed linear regression adjusting for age, age^2^, sex, and height, using rank-based inverse normal transforms, adjusting for population substructure using principal components or linear mixed models, and performing separate analyses for ever- and never-smokers or using a covariate for smoking (for studies of related subjects). Genomic control was applied to individual studies, and results were combined using fixed-effects meta-analysis^24^.

### Genome-wide association analysis

In UK Biobank, we performed logistic regression of COPD, adjusting for age, sex, genotyping array, smoking pack-years, ever smoking status, and principal components of genetic ancestry. Association analysis was done using PLINK 2.0 alpha^79^ (downloaded on December 11, 2017) with Firth-fallback settings, using Firth regression when quasi-complete separation or regular-logistic-regression convergence failure occurred. We performed a fixed-effects meta-analysis of all ICGC cohorts and UK Biobank using METAL (version 2010-08-01)^80^. We assessed population substructure and cryptic relatedness by linkage disequilibrium (LD) score regression intercept^81^. We defined a genetic locus using a 2-Mb window (+/−1 Mb) around a lead variant, with conditional analyses as described below.

To maximize our power to identify existing and discover new loci, we examined all loci at the genome-wide significance value of P < 5 × 10^−8^. We first characterized loci as being previously described (evidence of prior association with lung function^13,14,23,15–22^ or COPD^5,10–12^) or novel. We defined previously reported signals if they were in the same LD block in Europeans^82^ and in at least moderate LD (r^2^>= 0·2). For novel loci we attempted replication through association of each lead variant with either FEV_1_ or FEV_1_/FVC ratio in SpiroMeta, using one-sided p values with Bonferroni correction for the number of novel loci examined. Novel loci failing to meet a Bonferroni-corrected p value were assessed for nominal significance (one-sided p < 0·05) or directional consistence with FEV_1_ and FEV_1_/FVC ratio in SpiroMeta.

Cigarette smoking is the major environmental risk factor for COPD and genetic loci associated with cigarette smoking have been reported^5,83^. While we adjusted for cigarette smoking in our analysis, we further examined these effects by additionally testing for association of each locus with cigarette smoking and by looking at two separate analyses of ever- and never- smokers in UK Biobank.

### Identification of independent associations at genome-wide significant loci

We identified specific independent associations at genome-wide significant loci using GCTA-COJO^84^. This method utilizes an approximate conditional and joint analysis approach requiring summary statistics and representative LD information. As the UK Biobank provided the predominant sample, we used 10,000 randomly drawn unrelated individuals from this discovery dataset as a LD reference sample. We scaled genome-wide significance to a 2-Mb region, resulting in a locus-wide significant threshold of 8 × 10^−5^, or 2 × 10^−6^ for variants in the major histocompatibility complex (MHC) region (chr6:28477797-33448354 in hg19^85^). We created regional association plots via LocusZoom using 1000 Genomes EUR reference data (Nov2014 release)^86^.

### Identification and prioritization of tissues and cell types, candidate variants, genes, and pathways Identification of enriched tissues and specific cell types

We used LD Score Regression (LDSC)^87^ to estimate the enrichment of functional annotation in disease heritability. We utilized LDSC baseline models (e.g., conserved region, promoter flanking region), tissue-specific annotations from the Roadmap Epigenomics Program^28^ and GenoSkyline^27^. We also used SNPsea^29^ to estimate the enrichment of specific cell types from genome-wide significant associations using gene expression data (**Supplementary Methods**).For SNPsea, we used a single-cell RNA-seq dataset from the study of idiopathic pulmonary fibrosis (540 cells from 6 IPF lung samples and 3 control tissues, available at Gene Expression Omnibus as GSE86618)^30^ (**Supplementary Methods**).

### Fine-mapping of independent association signals at genome-wide significant loci

We used Bayesian fine-mapping at each locus to identify the credible set: the set of variants with a 99% probability of containing a causal variant. Briefly, for each genome-wide significant loci we calculated approximate Bayes factors^31^ of association. We then selected variants in each locus, so that their cumulative posterior probability was equal or greater than 0·99 using an unsealed variance. At loci with multiple independent associations, we used statistics from approximate conditional analysis with GCTA software on each index variant adjusting for other independent variants in the loci. Otherwise, we used unconditioned statistics from our meta-analysis. We characterized variant effects in credible sets using variant annotations from Ensembl Variant Effect Predictor^88^.

### Identification of target genes

We used several computational approaches with corresponding available datasets to identify target genes in genome-wide significant loci. We used two methods that utilized gene expression data: 1) S-PrediXcan and 2) DEPICT. We used S-PrediXcan^89^ to identify genes with genetically regulated expression associated with COPD. We used data from the Lung-eQTL consortium^90,91^ (1,038 lung tissue samples) as an eQTL and gene expression reference database. S-PrediXcan is the extension of PrediXcan^92^ that test for association between a trait and imputed gene expression using summary statistics. Here, we performed S-PrediXcan using models for protein-coding genes +/− 1 Mb from top-associated variants at genome-wide significant loci. We used DEPICT (Data-driven Expression Prioritized Integration for Complex Traits)^93^ to prioritize genes from reconstituted gene sets.

We also used additional information on gene regulation, including epigenetic data: 1) regulatory fine mapping, 2) mQTL, and 3) chromosome conformation capture. We used regulatory fine mapping (regfm^94^) to overlap 99% credible interval (CI) variants at each GWAS locus with open chromatin regions based on DNAse hypersensitivity sites (DHS). DHS cluster accessibility state was then associated with gene expression levels (for 13,771 genes) from 22 tissues in the Roadmap Epigenomics Project^95^. Using both the 99% CI and DHS overlap, as well as the DHS state and transcript level association, regfm calculates a posterior probability of association of each gene +/− 1 Mb of the lead SNP at each GWAS locus. We also searched for overlapping methylation quantitative trait loci (mQTL) data from lung tissue, as recently described^96^. To determine whether these signals co-localized (rather than being related due to linkage disequilibrium), we performed colocalization analysis between our GWAS and mQTL in genome-wide significant loci using eCAVIAR^97^ (eQTL and GWAS CAusal Variants Identification in Associated Regions, **Supplementary Methods**). We also sought information from publicly available chromosome conformation capture data^98^. We obtained statistics of high-confidence chromatin interaction in fetal lung fibroblast cell line (IMR90) and human lung tissue^98^ through HUGIn^99^ (Hi-C Unifying Genomic Interrogator). We anchored long range chromatin interactions on top associated variants and computed statistical significance of contact at each locus. We retained only the strongest associations (i.e., smallest P value) for each cell line/primary cell in the analysis.

Finally, we searched for signals from non-synonymous variants. We identified coding variants present in the credible set in the GWAS with a high posterior probability. We also searched for rare coding variants, based on exome sequencing results in the COPDGene, Boston Early-Onset COPD, and International COPD Genetics Network studies. In brief, we performed exome sequencing on 485 severe COPD cases and 504 smoking resistant controls from the COPDGene study and 1,554 subjects ascertained through 631 probands with severe COPD from the Boston Early-Onset COPD study (BEOCOPD) and the ICGN study. We performed single-variant analyses using Firth and efficient resampling methods (SKAT R package^100^) for the COPDGene data (case-control) and generalized linear mixed models (GMMAT) for the BEOCOPD-ICGN data (using lung function). Gene-based analyses were conducted using Burden, SKAT, and SKAT-O tests with asymptotic and efficient resampling methods (SKAT package) combined with Fishers method for the COPDGene data, and using SKAT-0 tests (MONSTER) for the BEOCOPD-ICGN data. Two variant-filtering criteria were considered: deleterious variants (predicted by FATHMM) with MAF < 0·01, and functional variants (moderate effect predicted by SNPEff) with MAF < 0·05. Gene-based segregation test (GESE) was also applied to the ultra-rare (MAF < 0·1%) and loss-of-function variants in the BEOCOPD-ICGN data on the severe COPD affection status.

For each dataset described above, we used Bonferroni-corrected P values, or a fixed posterior probability threshold to determine target genes at each locus. To correct for possible number of genes in each locus, we obtained a list of protein-coding genes +/−1 Mb from a top associated variant from BioMart^88^. For each locus, we used a 5% Bonferroni-corrected threshold (i.e., P < 0·05 divided by number of genes at that locus) to determine significance for 4 data types: gene expression data, chromatin conformation capture data, co-regulation of gene expression, and exome sequencing results. For two remaining datasets, we used a fixed posterior probability (of gene association with a GWAS locus) threshold of 0·1 for regfm and eCAVIAR. We considered genes that were implicated by gene expression or >= 2 combination of other datasets (e.g., methylation and chromatin conformation capture data) as target genes.

### Identification of pathways

To identify enriched pathways in COPD-associated loci, we performed gene-set enrichment analysis using the “reconstituted” genes sets from DEPICT, as described above^93^. We defined significant gene sets using false discovery rate (FDR) < 5%.

### Effects on COPD-related and other phenotypes

COPD is a complex and heterogeneous disorder, comprised of different biologic processes and specific phenotypic effects. In addition, many loci discovered by GWAS have pleiotropic effects. To identify these effects, we performed analyses of a) identification of overlapping genetic loci between related disorders (asthma and pulmonary fibrosis) b) genetic association studies of our genome-wide significant findings using COPD-related phenotypes, including a cluster analysis to identify groups of variants that may be acting via similar mechanisms; c) look up of top variants in prior COPD-related quantitative computed tomography (CT) imaging feature GWAS, and d) look up of associations with other diseases/traits using GWAS Catalog.

To identify overlapping loci between COPD and other respiratory disorders, we used gwas-pw^101^ to perform pairwise analysis of GWAS. This method searches for shared genomic segments^82^ using adaptive significance threshold, allowing detection of sub genome-wide significant loci. We identified shared segments or variants using posterior probability of colocalizing greater than 0·9^101^. We obtained GWAS summary statistics from two previous studies of pulmonary fibrosis^39^ and GWAS of asthma in Europeans^38^. To assess heterogenous effects of COPD susceptibility loci on COPD-related features (phenotypes), we evaluated associations of our genome-wide significant SNPs with 121 detailed phenotypes (e.g., lung function, computed tomography-derived metrics, biomarkers, and comorbidities) available in 6,760 COPDGene non-Hispanic whites. We calculated Z-scores for each SNP-phenotype combination relative to the COPD risk allele to create a SNP by phenotype Z-score matrix. We tested each COPD-related phenotype with at least one nominally significant association with one of our genome-wide significant COPD SNPs, leaving us with 107 phenotypes. We then oriented all Z-scores to be positive (based on sign of median Z score) in association with each phenotype to avoid clustering based on direction of association. To avoid clustering phenotypes only by strength of association with SNPs, we scaled Z-scores within each phenotype by subtracting mean Z-scores and dividing by the standard deviation of Z-scores within each phenotype. We then scaled Z-scores across SNPs to circumvent clustering of SNPs according only to relative strength of association with phenotypes. We then performed hierarchical clustering of the scaled Z-scores of associations between SNPs and phenotypes to identify clusters of SNPs and phenotypes for all 107 phenotypes as well as in the subset of 26 quantitative imaging phenotypes. We identified optimal number of clusters using the Calinski index^102,103^. To identify features that independently predict cluster membership, we fitted a logistic regression model via penalized maximum likelihood using the glmnet package^104^. We determined optimal regularization parameters using 10-fold cross validation. We further examined top variant associations with COPD-related traits through a look-up of top variants in a prior GWAS of 12,031 subjects with quantitative emphysema and airway CT features^105^. To examine overlap of our COPD results with other traits, we downloaded genome-wide significant associations from the GWAS Catalog^37,106^ (P < 5 × 10^8^). Between a pair of COPD- and trait-associated variants within the same LD block in Europeans^82^, we computed the LD using the European ancestry panel and considered the overlap if variants were in at least in moderate LD (r^2^ >= 0·2).

## Tables

Table 1 Meta-analysis results showing 35 loci novel for COPD and lung function (see the Excel file)

## Acknowledgements

Please refer to the **Supplementary Note** for full acknowledgements.

## Author contributions

P.S. contributed to the study concept and design, data analysis, and manuscript writing. D.P., B.D.H., M.H.C. contributed to the study concept and design, data analysis, statistical support, and manuscript writing. A.B.W., K.d.J., S.J.L., D.P.S. contributed to the study concept and design and data analysis. P.B., R.G.B., J.D.C., A.G., D.A.M., G.T.O., S.I.R., D.A.S., R.T.-S., Y.T., E.K.S. contributed to the study concept and design and data collection. T.H.B., J.E.H. contributed to the study concept and design and to statistical support. I.P.H., H.M.B., L.V.W., M.D.T. contributed to the study concept and design. All authors, including those whose initials are not listed above, contributed to the critical review and editing of the manuscript and approved the final version of the manuscript.

## Supplementary Figures

**Supplementary Figure 1 Forest plots for 82 genome-wide significant associations**

**Supplementary Figure 2 Regional association plots for 82 genome-wide significant associations**

**Supplementary Figure 3 Enrichment of genes specifically expressed in respiratory cell types**

**Supplementary Figure 4 Number of variants in 99% credible sets**

**Supplementary Figure 5 Heatmap of associations of 82 index variants and phenotypes in COPDGene**

**Supplementary Figure 6 Associations of index variants and traits in NHGRI-EBI GWAS Catalog**

**Supplementary Figure 7 Comparison of odds ratios (OR) including and excluding individuals with asthma of 82 genome-wide significant variants**

### Supplementary Tables

See the Excel file.

**Supplementary Table 1 Cohort baseline characteristics in COPD cases and controls**

**Supplementary Table 2 Meta-analysis results showing 47 previously reported loci for COPD or lung function**

**Supplementary Table 3 Multiple independent associations within the same 2-Mb window identified using approximate conditional and joint analysis**

**Supplementary Table 4 Heritability enrichment in cell-type specific epigenomic mark from Roadmap Epigenomic Project**

**Supplementary Table 5 Genes specifically expressed in genome-wide significant loci**

**Supplementary Table 6 Functional annotation of variants with posterior probability of association greater than 0.6**

**Supplementary Table 7 Candidate target genes**

**Supplementary Table 8 Gene sets significantly enriched at FDR < 0.05 using DEPICT Supplementary Table 9 Overlapping loci between COPD with asthma and pulmonary fibrosis**

**Supplementary Table 10 Lookup associations for quantitative computed tomography (QCT) features and cluster membership**

**Supplementary Table 11 Association results from GWAS catalog**

**Supplementary Table 12 Genetic correlation between COPD and other traits/diseases**

